# Drug Combination Antagonism and Single Agent Dominance Result from Differences in Death Activation Kinetics

**DOI:** 10.1101/805093

**Authors:** Ryan Richards, Hannah R. Schwartz, Mariah S. Stewart, Peter Cruz-Gordillo, Megan E. Honeywell, Anna J. Joyce, Benjamin D. Landry, Michael J. Lee

## Abstract

Therapeutic regimens for cancer generally involve drugs used in combinations. Most prior work has focused on identifying and understanding synergistic drug-drug interactions; however, understanding sources of antagonistic interactions remains an important and understudied issue. To enrich for antagonistic interactions and reveal common features of these drug combinations, we screened all pairwise combinations of drugs characterized as canonical activators of different forms of regulated cell death. We find that this network is strongly enriched for antagonistic interactions, and in particular, enriched for an extreme form of antagonism, which we call “single agent dominance”. Single agent dominance refers to antagonisms in which a two drug combination phenocopies one of the two agents. We find that dominance results from differences in the cell death onset time, with dominant drugs inducing death earlier and at faster rates than their suppressed counterparts. Finally, we explored the mechanisms by which parthanatotic agents dominate apoptotic agents, finding that dominance in this scenario is caused by mutually exclusive and conflicting use of PARP1. Taken together, our study reveals death activation kinetics as a predictive feature of antagonism, due to inhibitory crosstalk between cell death pathways.

## INTRODUCTION

Cancer therapies are often limited by issues such as acquired drug resistance and partial killing of a population of tumor cells^1, 2^. To combat these limitations, many efforts focus on the development of combination drug therapies^3–5^. Generally, most prior studies have focused on identifying combinations that produce synergistic drug-drug interactions. In contrast to expectations, however, recent reports have demonstrated that synergy is not generally observed in clinically efficacious drug combinations, which instead are typically additive^6^. Synergistic drug combinations tend to reinforce the killing that would be induced by one of the single agents within the combination, rather than facilitating the killing of new cells that would not be killed by either of the single agents alone^7, 8^. Furthermore, synergistic combinations may also favor the future evolution of resistant clones^9^. While these data may limit the value of identifying drug synergies, understanding the sources of drug-drug antagonism still remains an important issue. It stands to reason that antagonism – particularly very strong antagonism – may limit the efficacy of a drug combination. Predicting which drug combinations will result in antagonism is challenging due to the lack of transparent “rules” underlying this phenomenon, as well as the unpredictable and often genotype specific nature of drug-drug interactions^10^. Thus, an unmet need is the identification of robust guiding principles that can be used to more efficiently identify, predict, or even improve upon antagonistic drug-drug interactions.

In the absence of robust principles to enable prediction of non-additive drug interactions, a common approach is to screen drug combinations, prioritizing the testing of drugs that target proteins with complementary functions. Several recent studies have successfully used known or predicted network topologies to enrich for non-additive drug combinations^11–14^. Furthermore, network simulations have revealed topological features, such as negative feedback and mutual inhibition that may underlie the antagonism of drug combinations^15^. We envisioned that principles of drug-drug antagonism may emerge from studying drugs that target a network that was enriched for antagonistic interactions.

In recent years it has become clear that at least twelve mechanistically distinct forms of regulated cell death exist in mammalian cells^16^. Because these death pathways function in a mutually exclusive manner, we reasoned that drug combinations designed to co-activate multiple types of cell death may be enriched for antagonistic drug-drug interactions. Several lines of evidence already exist to suggest a degree of negative interaction and/or interdependent and mutually exclusive function among the various forms of cell death^17, 18^. For instance, activation of necroptosis requires inhibition of extrinsic apoptosis, due to cleavage of the pro-necroptotic protein RIPK1 by caspase-8^19^. Similarly, PARP1, the initiator of parthanatos, is cleaved by caspase-3, suggesting that activation of apoptosis also inhibits the cell’s ability to activate parthanatos^20^. Although most potential interactions between death pathways have not explored, a model is beginning to emerge from these studies that mutually exclusive activation of cell death pathways may be enforced through inhibitory crosstalk between death regulatory pathways^16^.

To identify a robust set of antagonistic drug-drug interactions, we tested all pairwise combinations of the canonical activators for different cell death subtypes. We find that drug combinations comprised of cell death targeting drugs are enriched for drug antagonism, and in particular, strongly enriched for an extreme form of antagonism that we call “single agent dominance” (SAD). In SAD combinations, the two drug combination phenocopies one of the two single drugs. Importantly, this occurs even when the dominant drug is the less efficacious of the two compounds. Using multivariate statistical modeling we find that a key feature of SAD combinations is a large discrepancy in the timing of cell death onset, with faster acting drugs suppressing slower acting drugs, leading to strong antagonism. These antagonistic phenotypes could be relieved by temporally phasing drug addition to promote synchronized co-activation of multiple death pathways. Finally, we explore the molecular mechanisms of SAD combinations involving apoptotic and parthanatotic agents, finding that mutually exclusive but conflicting use of PARP1 drives dominance in these scenarios. Taken together, these findings highlight that the interconnected nature of cell death processes causes unexpected behaviors when these pathways are co-activated. Furthermore, we find that the rate of activation is a feature of antagonistic drug responses, with a previously unappreciated role in dictating functional interactions between cell death processes.

## RESULTS

### Optimization of a high-throughput assay for monitoring cell death kinetics

Drugs that induce different forms of cell death vary substantially in terms of their potency and their rates of activation^21^. Thus, to evaluate drug combinations comprised of apoptotic and non-apoptotic agents, we first optimized an assay that could be performed in a high-throughput scale while also retaining accurate analysis of death activation kinetics. Recently, methods have been developed for high-throughput measurement of cell death using SYTOX green in a fluorescence plate reader^17^. SYTOX green is a cell impermeable dye that fluoresces when bound to double stranded DNA. Furthermore, SYTOX fluorescence is specific to cell death, but largely agnostic to the mechanism by which cells die^22^. We made experimental and computational modifications to prior methods to enable an accurate inference of the numbers of live cells, dead cells, and total cells throughout the assay, rather than focusing only on the dead cells (Fig. 1a). Quantification of drug-induced death kinetics requires an accurate quantification of both live cells and dead cells over time, as inferences built from only one or the other of these measurements can be misleading^21^. To gain these insights, we measured of the initial lethal fraction prior to drug addition (e.g. percentage of dead cells relative to total cells), and at the assay end point (Fig. 1a). To determine the lethal fraction at intermediate time points throughout the experiment, experimentally measured dead cell numbers (raw SYTOX values) at intermediate times were compared to computationally inferred total cell numbers at these time points (Fig. 1a and Supplementary Fig. 1). This procedure enables the quantification of drug-induced changes in growth rate, death rate, and lethal fractions over time, including an analytical estimation of death onset time (Fig. 1b and Supplementary Fig. 1). Furthermore, all of these measurements are computed from a single assay, and without the need for any specialized equipment. To evaluate the sensitivity of our assay we tested manual two-fold cell dilutions over a wide range of cell concentrations. We found that SYTOX green fluorescence of U2OS cells was linearly correlated with cell number from approximately 50 – 20,000 cells per well, allowing accurate quantification of death rates even as low as 1% above background (Fig. 1c). To determine the accuracy of our assay – and in particular the accuracy of our death rate estimates – we compared our approach to STACK, an automated-microscopy-based approach that enables direct measurement of death kinetics by simultaneously measuring live and dead cells^21^. Overall, we found a strong correlation between our computationally inferred death rates and those observed using direct measurement (Fig. 1e).

**Figure 1.**
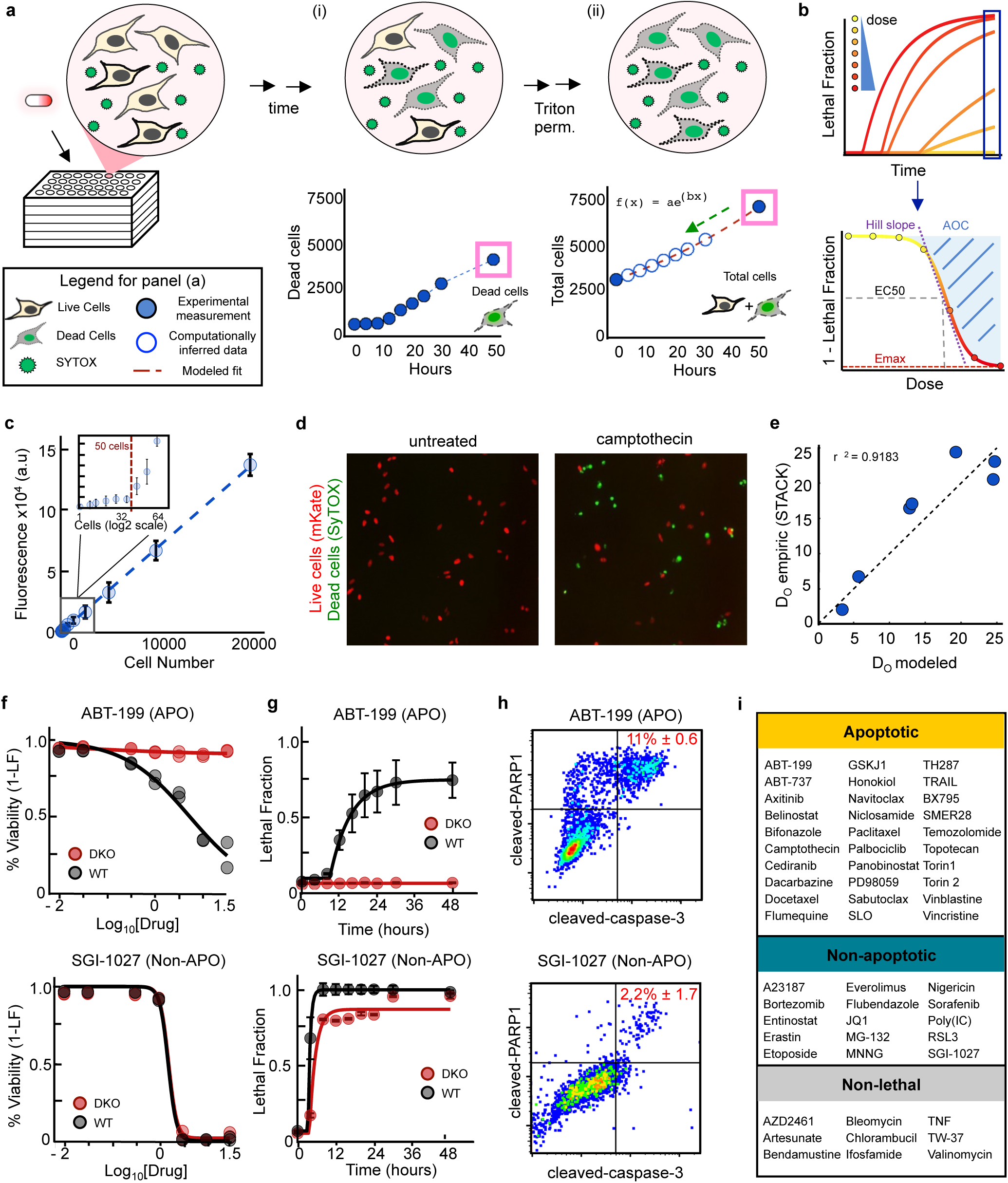
Characterization of drug mechanisms of action using a high-throughput assay to quantify cell death kinetics. **(a)** Schematic of SYTOX assay. Cells are treated with drug in the presence of 5 µM SYTOX Green to count dead cells. (i) Fluorescence measured every 4 hours for 48 hours which is linearly proportional to dead cell number. (ii) After triton permeabilization total cell counts can be determined at 0 hour and 48 hour time points (filled blue circles). Total cell counts can be modeled to an exponential fit (red dashed line) to calculate total cell counts at measured time points (unfilled blue circles). **(b)** Modeled total cell counts were used to infer the Lethal Fraction (LF) at all time points throughout the assay. LF was fit to a kinetic model to quantify death onset (D_O_) and rate (D_R_) following cell permeabilization to count total cells. See also Supplementary Fig. 1. Percent viability at 48 hours was fit to a 4 parameter logistic regression to quantify pharmacological parameters. **(c)** SYTOX plate reader sensitivity. Known cell counts from serial dilution or cell sorting were lysed in 5 µM SYTOX Green to test the linearity and detection limit of the assay. **(d-e)** Comparison of computational estimates of drug activation rate to empiric measurements using the STACK protocol. **(d)** Example image from cells treated with camptothecin (3.16 µM) or untreated. **(e)** Correlation between death onset time measurements calculated from imaging or plate reader based estimates. **(f-i)** Classification of drug mechanism of cell killing. For panels (f-h), examples of an apoptotic (top) or non-apoptotic (bottom) drug are shown. Drugs were tested in U2OS (WT) or U2OS-BAX/BAK^-/-^ (DKO) cells at 7 doses over 48 hours. **(f)** Percent viability across varied doses. **(g)** Lethal Fraction kinetics at the highest drug concentration, measured over time **(h)** Apoptotic activity quantified by flow cytometry. Cleaved-PARP1/cleaved-Casp3 double positivity was quantified 24hr post-treatment. Percentages are the mean +/- standard deviation. **(i)** List of apoptotic and mixed/non-apoptotic drugs. See also Supplementary Fig 2-4. For panels **(c)** and **(g)** error bars represent the standard deviation among replicates.

Using our assay, we next characterized a panel of 54 drugs that are reported to kill cells using different forms of regulated cell death^16, 23–30^. Drug responses were screened in U2OS cells, a cancer cell line with wild-type p53, which respond well to a diverse array of cell death inducing agents. Due to limited availability of markers to measure activation of some forms of death, we focused on a simplified classification scheme, categorizing drugs as non-lethal, apoptotic, or non-apoptotic. Non-lethal compounds were those whose effects were exclusively due to modulation of growth rate rather than through cell killing (Supplementary Fig. 2-4). To distinguish between apoptotic and non-apoptotic drugs, we scored the degree to which the observed drug response was modulated in a BAX/BAK double knockout (DKO) genetic background relative to the wild-type (WT) parent U2OS cells (Fig. 1f-g). BAX and BAK are members of the BCL2 family of proteins and are pore forming proteins required for mitochondrial outer membrane permeabilization and subsequent apoptosis^31^. Drug classifications based on relative DKO vs. WT sensitivity were in agreement with categorization based on caspase-3 cleavage, a marker of activation of apoptosis (Fig. 1h). Overall, our classification strategy identified 9 non-lethal compounds, 30 apoptotic drugs, and 15 non-apoptotic drugs (Fig. 1f-i and Supplementary Fig. 2-4).

### Co-activation of apoptotic and non-apoptotic death is enriched for antagonism and single agent dominance

The different subtypes of regulated cell death are activated in a mutually exclusive manner, due to inhibitory interactions between the death pathways. These features have been found to be common among antagonistic drug-drug interactions^15^. Thus, we next aimed to test if co-activation of multiple death pathways would enrich for drug antagonism. Our strategy was to test all pairwise combinations of the 54 cell death-activating drugs that we evaluated as single agents. All single drugs and drug combinations were tested across 7 doses at a fixed dose ratio, and responses were measured every 4 hours over 48 hours (Fig. 2a-b). Biological replicates within the screen were highly correlated, suggesting a high degree of reproducibility within our assay (r2 > 0.98, Fig. 2c). To score drug-drug interactions, we compared the observed responses to predicted responses given independent drug action (Fig. 2b). Overall we found 265 drug combinations that resulted in synergy, and 560 combinations that resulted in antagonism (Fig. 2d). These data were highly reproducible upon retesting with other assays (r2 > 0.8, Fig. 2e-f, and Supplementary Fig. 1d).

**Figure 2.**
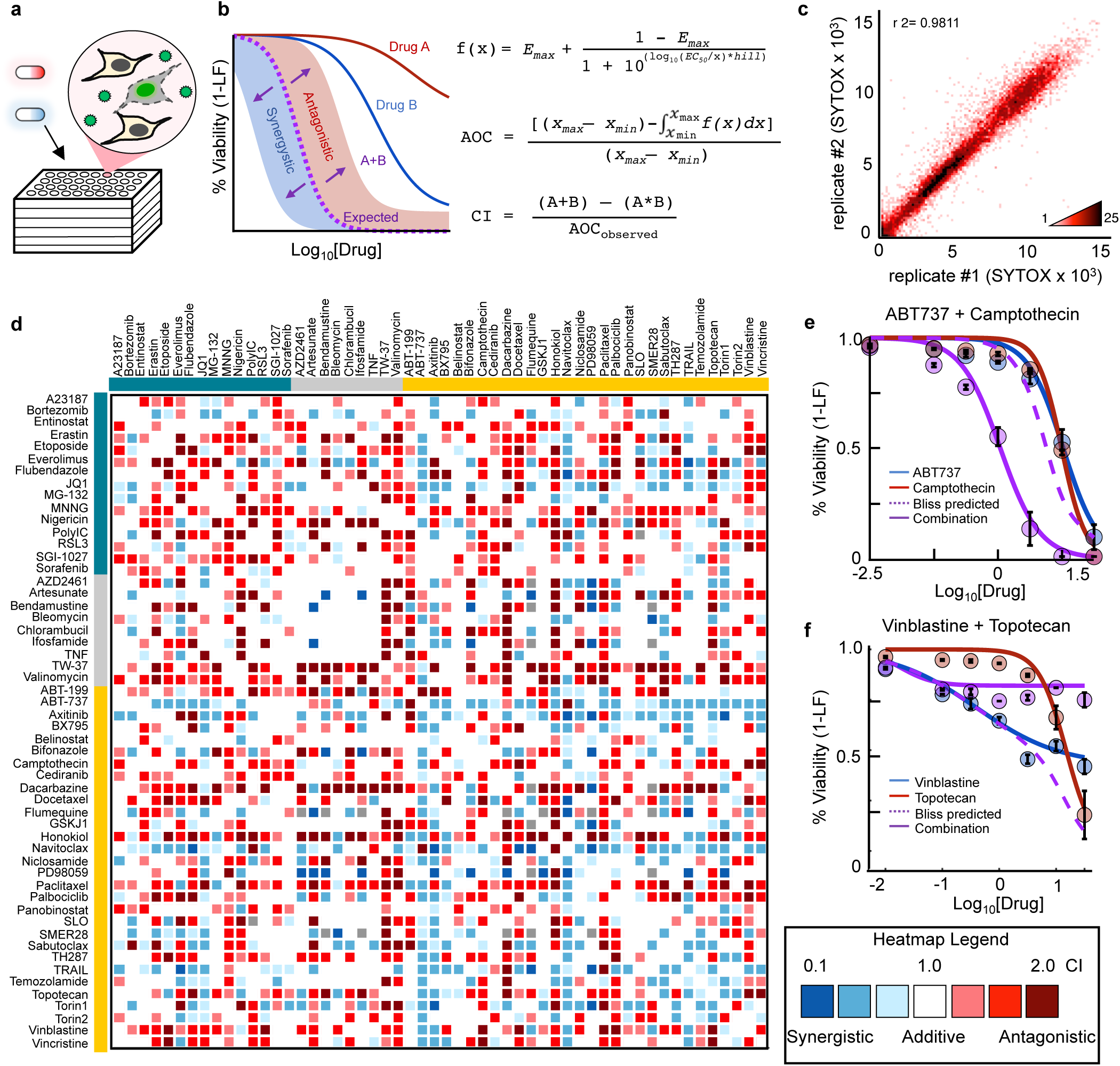
Combination drug screen to evaluate co-activation of apoptotic and non-apoptotic death pathways. **(a)** SYTOX Green assay for drug combinations. Drugs applied at fixed 1:1 dose ratio, and death was measured via SYTOX fluorescence over time. **(b)** Combination Index (CI) calculation. Equations shown for calculating dose curves, Area Over the Curve (AOC), and CI. CI based on expected AOC of the drug combination given individual drug responses (according to Bliss Independence) compared to the observed combination AOC. **(c)** Density plot of biological replicates from the drug combination screen. Pearson correlation coefficient shown. **(d)** Drug combination screen. Heatmap of all combination indices determined from screening 54×54 drug combinations. Colored bars depict drug class: Teal – Non-apoptotic; grey – non-lethal; orange – apoptotic. **(e-f)** Examples of known synergistic/antagonistic drug combinations. **(e)** ABT737 (blue) and camptothecin (red). **(f)** Vinblastine (blue) and topotecan (red). For **(e)** and **(f)** purple – observed combination; dashed purple – predicted combination assuming bliss independence. Data are mean +/- standard deviation.

Overall, in our “all-by-all” combinatorial drug screen nearly 60% of all combinations tested resulted in non-additive responses, with approximately 40% of all combinations resulting in antagonism (Fig. 3a-b). By comparison, other large-scale combination drug screens comprised of “random” drug combinations have identified antagonism at a frequency of 2-15% (Fig. 3a-b)^32, 33^. Furthermore, we found a strong enrichment for antagonism specifically with drug combinations that involved two different classes of compounds (p = 0.03, Fishers exact). Thus, as expected, our combination drug screen of cell death activating drugs is strongly enriched for antagonistic drug-drug interactions.

**Figure 3.**
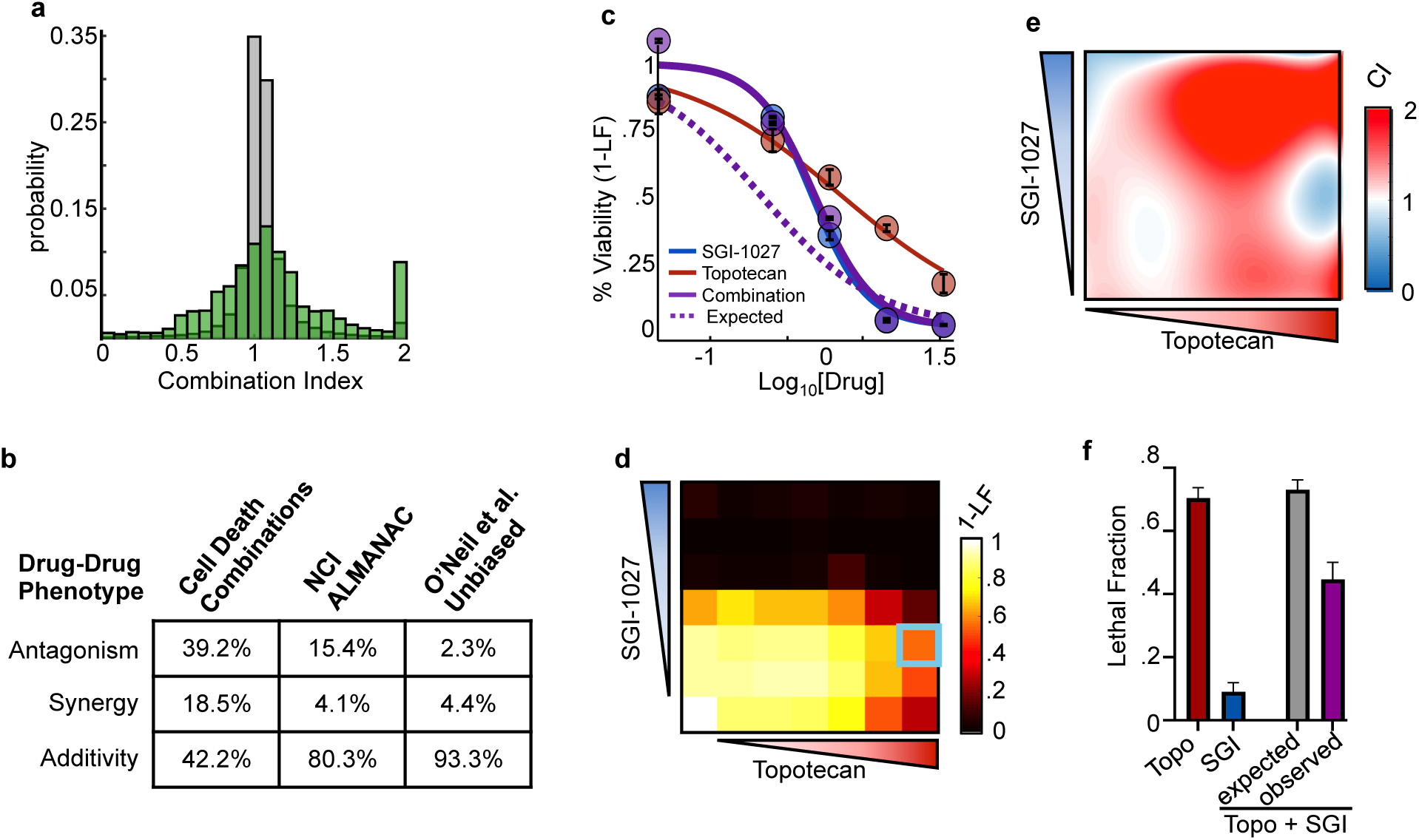
Combinations of apoptotic and non-apoptotic cell death drugs are enriched for antagonism and single agent dominance. **(a)** Histogram of combination indices (CI). CIs determined from NCI ALMANAC, and the present screen. Grey – NCI ALMANAC data; Green – Cell death screen. **(b)** Summary statistics comparison of antagonism, synergy, and additivity for drug combination screens. **(c)** Example of a SAD combination. Drugs given in 1:1 dose ratios and measured using SYTOX at 48 hours. Data are mean +/- standard deviation. Blue – SGI-1027; Red – Topotecan; Purple – Combination; Dashed purple – expected additivity. **(d)** Percent viability response matrix for all-by-all dose combinations. The bounded area represents the doses shown in **(f)** and depict suppression of lethal doses of topotecan by non-lethal doses of SGI-1027. **(e)** Combination indices for all-by-all dose combinations, calculated at each dose using Bliss independence. **(f)** Lethal Fraction from 10 µM Topotecan and 0.3 µM SGI-1027 treatment. Data are mean +/- standard deviation.

To more deeply analyze our data, we decided to focus on a large set of very strong antagonistic drug-drug interactions that we uncovered (Fig. 3a). While, nearly 40% of all combinations resulted in antagonism, many of these antagonistic responses were an extreme form in which the two drug combination phenocopied one of the two single agents (Fig. 3c). We call this response “single agent dominance” (SAD). The characteristic feature of SAD combinations is that an otherwise efficacious drug becomes fully suppressed by a second agent. For example, both SGI-1027, a DNA methyltransferase inhibitor that induces non-apoptotic death, and topotecan, a Topo I inhibitor that induces apoptosis, produced strong killing in U2OS cells when applied as single agents (Fig. 3c). The combination of these two agents, however, resulted in precisely the same response as SGI-1027 alone (Fig. 3c). One relatively trivial explanation of this phenotype could be that very potent drugs tend to dominate simply because no additional cells remain to be killed (e.g. maximum potency was achieved by a single drug). To test this idea directly we tested all dose-dose combinations (e.g. rather than fixed dose ratios), to determine if low concentrations of a dominant drug (e.g. SGI-1027) would suppress high concentrations of a suppressed drug (e.g. topotecan). We found that even non-efficacious concentrations of SGI-1027 were sufficient to block death induced by high concentrations of topotecan (Fig. 3d-f). In fact, nearly all combinations of SGI-1027 and topotecan resulted in antagonism, regardless of the doses used (Fig. 3f). Overall, we identified 179 SAD combinations, which were 36% of all antagonistic drug-drug interactions.

### The rate of drug-induced cell death is a key determinant of drug dominance

Considering the prominence of the SAD phenotype in our data, we next sought to gain a deeper understanding of the determinants of SAD antagonism by identifying features that were enriched within SAD combinations. In order to identify features of SAD combinations that might be helpful in predicting when/where these phenotypes will occur, we performed a multivariate analysis on the kinetic and pharmacological data generated by our SYTOX based cell death assay. To reduce the dimensionality of our data, we used principal component analysis (PCA). Using PCA, 6 principal components were identified that captured over 85% of the variation in drug pharmaco-kinetics (Fig. 4a). Projection of the data onto PC1 and PC2 revealed a clear enrichment of SAD combinations for positivity on PC1, which captured 23.5% of the overall variation within our dataset (Fig. 4b-c). To determine which other PCs were capturing information related to SAD combinations, we scored the statistical enrichment for SAD combinations on each PC. SAD combinations were also enriched on PC2, which captured 19% of the drug response variation, and to a lesser extent on PC3 (Fig. 4d).

**Figure 4.**
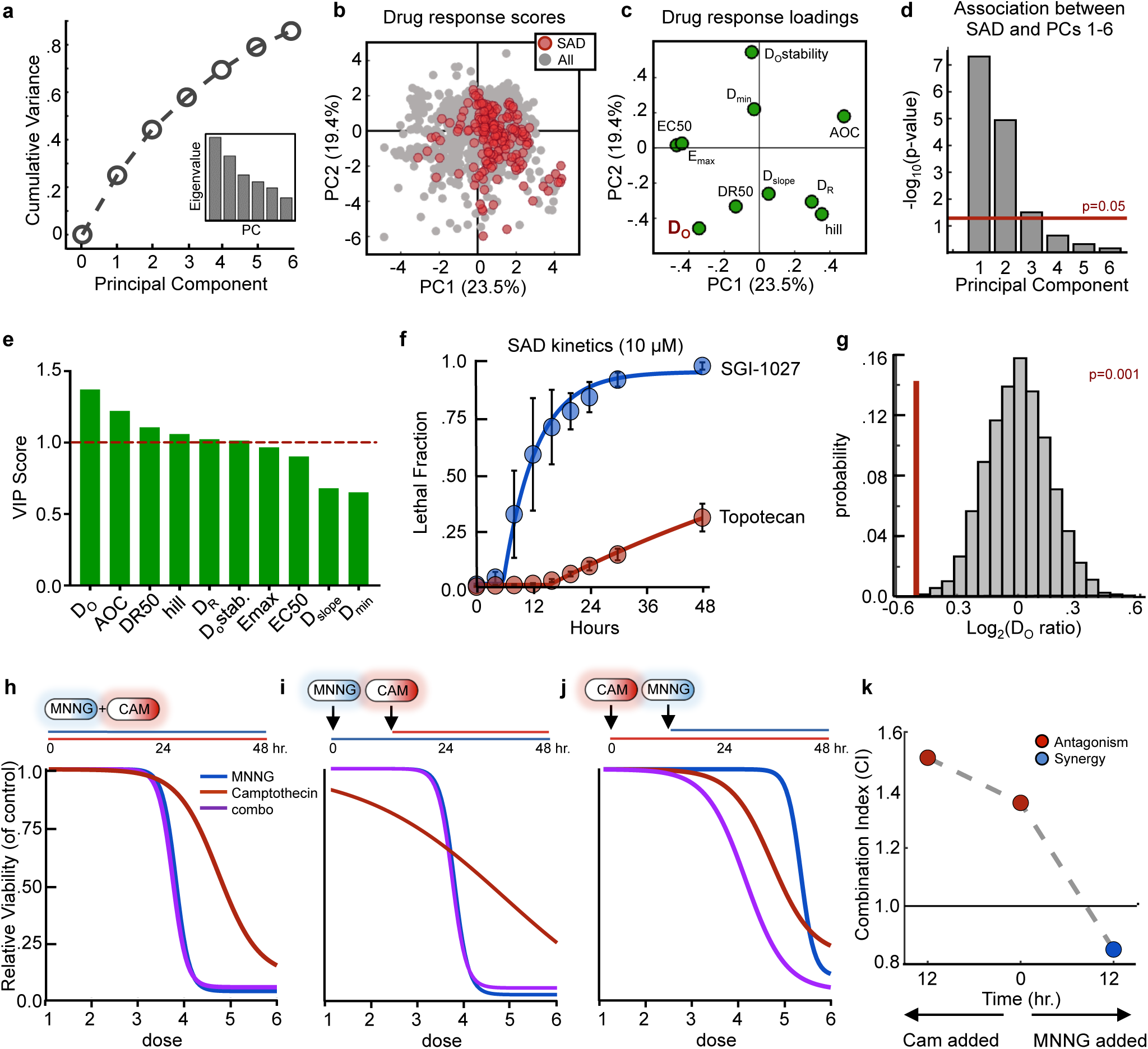
Statistical modeling reveals drug activation kinetics as a key determinant of SAD combinations. **(a)** Cumulative variation captured by PC1-6. Inset – Eigenvalues for PC1-6. **(b)** PCA scores plot (PCs 1 and 2) for all single drugs and combinations from the cell death screen. SAD combinations are colored red while all other combinations/drugs are colored grey. **(c)** Loadings plot for PCs 1 and 2. PCA was performed using 10 input variables consisting of kinetic and pharmacological parameters. **(d)** Enrichment of SAD for PCs 1-6. Enrichment for each PC was calculated using a one-tailed Fisher’s exact test. The red line depicts a 0.05 statistical cutoff. **(e)** Ranked VIP scores of PCA input variables. The red dashed line corresponds to a weighted importance cutoff of 1.0. **(f)** LF kinetics for an example SAD combination. Blue – dominant drug; red – suppressed drug. Data are mean ± S.D. **(g)** Distribution of average onset ratios for all combinations (grey) or SAD combinations (red). **(h-j)** Dose response curves of MNNG and camptothecin (CAM) combinations added simultaneously (h) or sequentially (i and j). Blue – MNNG; Red – CAM; Purple – combination. **(k)** Combination indices over time for MNNG + CAM. Red – antagonistic; blue – synergistic.

Considering the strong enrichment for SAD combinations on this subset of PCs, we next sought to determine what information about the drug response was being captured on PCs that are enriched for SAD combinations. To answer this question, we computed the Variable Importance in Projection, also called the VIP Score, which is a weighted sum of the relative magnitude of the regression coefficients for each input variable (i.e. a description of the importance of each variable in defining these principal components)^34^. Based on VIP scores, the onset time of cell death (D_O_) stood out as being the most important variable associated with SAD combinations (Fig. 4e). To determine what aspect of death onset time was related to the SAD phenotype, we examined the kinetics of cell death for the individual drugs that make up SAD combinations. We noticed a strong discrepancy in the rates of activation of dominant and suppressed drugs. For instance, in combinations of SGI-1027 and topotecan – a combination in which SGI-1027 is dominant – SGI-1027 kills cells earlier and at a faster rate than topotecan (Fig. 4f). To determine if differences in rates of activation were a common feature of SAD combinations, we explored the kinetic ratios for all 179 SAD combinations compared to 10,000 iterations of 179 random drug combinations in our data. This analysis confirmed that dominant drugs activate significantly faster and kill significantly earlier than their suppressed counterparts (Fig. 4g).

We next tested if the strong statistical correlation between onset time asymmetry and drug dominance was indicative of a causative relationship. To address this question our strategy was to temporally stagger the addition of dominant drugs, such that their activity occurred concomitantly with their suppressed counterparts. We tested this concept with the combination of MNNG and camptothecin, which results in robust domination by MNNG (Fig. 4h). The addition of the suppressed drug later in time relative to the dominant drug did not affect overall drug response or the degree of MNNG dominance (Fig. 4i); however, drug-drug antagonism and drug dominance were relieved when drug addition was staggered such that the suppressed agent is active at the same time as the dominant drug (Fig. 4j-k). Thus, taken together these data suggest that faster acting drugs suppress slower acting drugs, leading to drug-drug antagonism and single agent dominance.

To determine the generalizability of these findings, we aimed to determine if differences in death onset time could be predictive of drug-drug antagonism and new SAD phenotypes in the absence of any other information about the drugs and their mechanisms of cell killing. We used publically available data on single drugs that have not yet been tested in combinations, but for which drug-induced death kinetics were available. Full kinetic data for drugs at varied doses were not available for any drugs; however, kinetic data were available for limited doses for a large panel of drugs from a recent study that explored the kinetics of cell death^21^. Our strategy to generate robust predictions from this sparse kinetic data was to incorporate the pharmacological and kinetic data for new drugs into our PCA model, along with publically available pharmacological (i.e. “dose response”) data for drug responses in U2OS cells^35^. These data were used to determine relative vector projections for new drugs in PC1 and PC2, which were associated with SAD combinations (Supplementary Fig. 5-6).

Using this publicly available data, we identified 78 combinations among 15 drugs whose relative single agent profiles in PC1 and PC2 were similar to SAD combinations, suggesting drug dominance. We validated these predictions using our SYTOX based death assay (Supplementary Fig. 6). In our validation screen, greater than 94% of the predicted SAD combinations resulted in drug antagonism, with 55% of these resulting in a SAD interaction (Supplementary Fig. 6f-h). Thus, even in the absence of any information about the drugs and their mechanisms of action, an analysis of single drug pharmaco-kinetic properties was sufficient to identify new SAD combinations.

### PARP1 dependent interactions between parthanatotic and apoptotic death mediate drug dominance

Our drug combination screen and subsequent computational analyses have revealed the overall prevalence and a general mechanism driving single agent dominance. Next we sought to gain a molecular understanding of these drug-drug interactions. Non-additivity is a feature often related to the underlying network structure of a drug’s target proteins^12^. Given the diversity of the drugs tested and the limited understanding of pathways that regulate most forms of non-apoptotic death, we focused on studying a SAD combination that featured drugs with known regulatory pathway interactions. For example, hyper-activation of PARP1 causes parthanatos, a form of non-apoptotic cell death^16, 36^. Conversely, inhibition of PARP1 causes apoptosis due to “PARP trapping” and stabilization of DNA lesions^37^. Furthermore, PARP1 itself is cleaved and inactivated by caspase-3^20^. These data suggest a negative interaction between apoptotic and parthanatotic pathways, possibly due to the conflicting use of PARP1. To test this, we first evaluated the pathway activation dynamics of parthanatotic and apoptotic drugs at the level of PARP1 activation or inhibition. MNNG exposure leads to a transient hyper-activation of PARP1, resulting in increased protein PARylation (Fig. 5a). Drug-induced PARP1 hyper-activation began minutes after drug exposure, and activity returned to baseline by roughly one hour following MNNG exposure. When treated with apoptotic drugs like camptothecin, PARP1 was not activated, but rather was cleaved and inactivated by caspase-3 (Fig. 5b). PARP1 cleavage did not occur until roughly 8-12 hours after camptothecin exposure (Fig. 5c). Thus, following exposure to MNNG or camptothecin, perturbations to PARP1 activity occur in distinct temporal windows, and in a mutually exclusive manner. When cells are exposed to both MNNG and camptothecin, PARP1 activation follows an MNNG-like dynamic pattern, consistent with the observed phenotypic dominance by MNNG over camptothecin (Fig. 5a-c).

**Figure 5.**
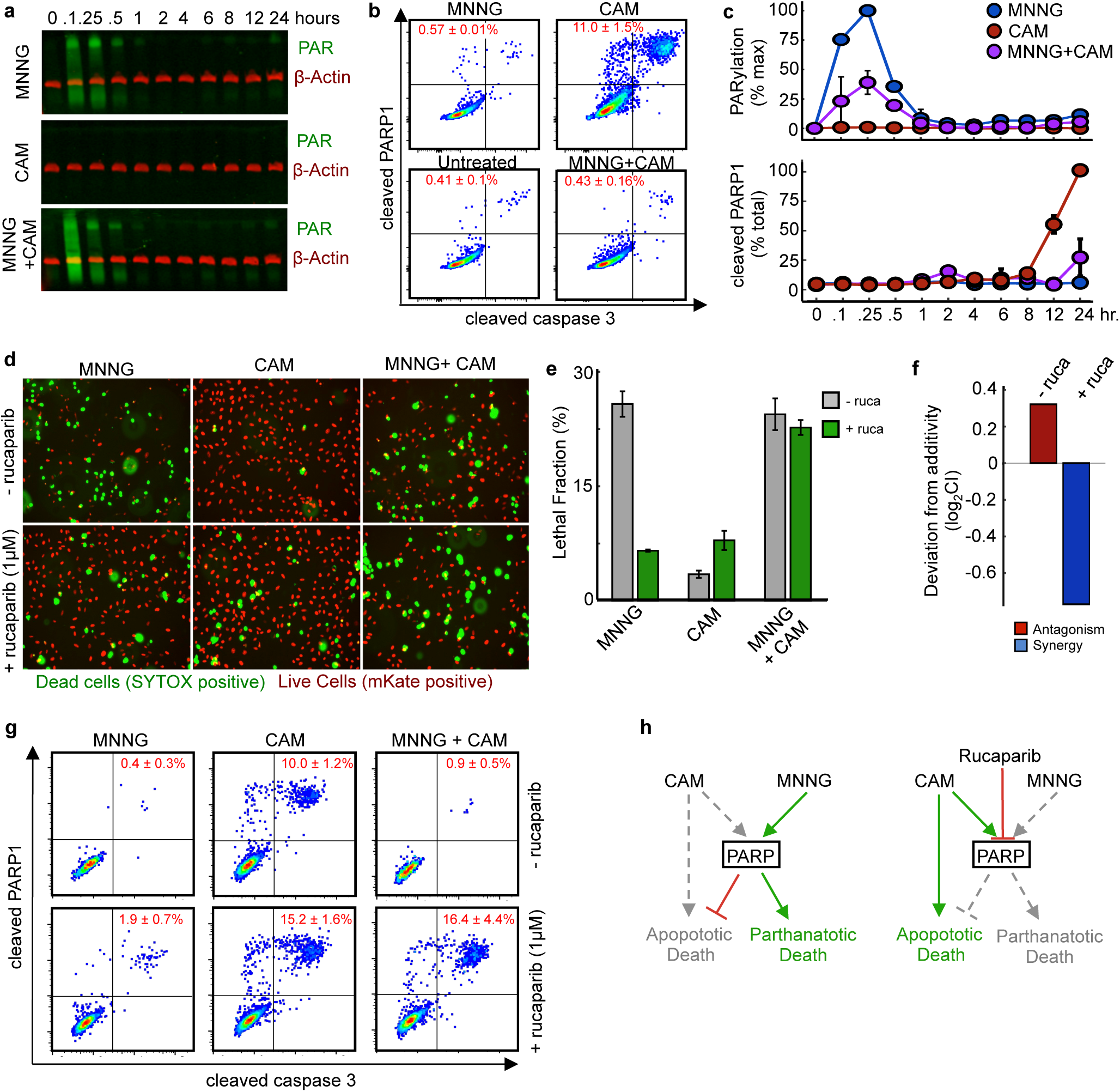
PARP1-dependent interactions mediate single agent dominance and choice between parthanatotic or apoptotic death. (a-c) Dynamics of drug-induced changes in PARP1 activity. Samples were treated with MNNG (31.6 µM), camptothecin (CAM, 3.16 µM), or MNNG+CAM for indicated times. **(a)** Western blot of total protein PARylation. Green – total PAR; red – b-actin. **(b)** Cleaved-PARP1 quantified by FACS (12 hr. treatment). Data are % cleaved PARP1 as mean +/- SD. **(c)** Quantification of PAR (top) and cleaved-PARP1 (bottom) activation dynamics. PAR quantified as % max signal. Cleaved PARP1 is shown as a percent of total cells. Values represent the mean +/- SD. **(d)** Quantitative live cell imaging using STACK assay. U2OS cells expressing nuclear localized mKate2 (red) were treated with MNNG, CAM, or MNNG+CAM with or without 1 µM rucaparib (ruca). SYTOX positive cells (green) are dead. Representative images from 48 hours post treatment. **(e)** Quantification of imaging in panel (d). Data are mean +/- SD of biological replicates. **(f)** Deviation from expectation. Expected combination response calculated using Bliss independence using the quantified LF values from (e). **(g)** Apoptotic activity of MNNG+CAM. Cleaved-PARP1/cleaved-caspase3 quantified 12 hours after treatment. Data are the mean +/- SD. **(h)** A model for a PARP mediated interaction between parthanatotic and apoptotic death leading to SAD.

Having determined that parthanatotic and apoptotic perturbations of PARP1 occur in a mutually exclusive manner, we next sought to determine if PARP1 is mechanistically involved in the MNNG-camptothecin SAD phenotype. To do so, we explored the fate of cells treated with MNNG and/or camptothecin in the presence or absence of a PARP1 inhibitor, rucaparib. Although PARP1 inhibition by rucaparib can itself induce apoptosis, we used rucaparib at a sub-lethal dose, which was sufficient for blocking PARP1 activity (Supplementary Fig. 7). Consistent with prior expectations, PARP1 inhibition largely blocked MNNG-mediated cell death, but enhanced camptothecin sensitivity (Fig. 5d-e). Interestingly, the MNNG dominance over camptothecin was lost when PARP1 was inhibited by rucaparib (Fig. 5e-f). Instead, in the presence of rucaparib, combinations of MNNG and camptothecin led to synergistically enhanced levels of cell death (Fig. 5f). Parthanatotic death, by definition, should not occur when PARP1 is inhibited^16^. Thus, we suspected that PARP1 inhibition changed not only the nature of the drug-drug interaction between MNNG and camptothecin, but also changed the mechanism of cell death induced by this combination. To test this, we began by evaluating markers of activation of apoptotic cell death. Indeed, using a flow cytometry based measurement of caspase-3 activation, we found that PARP1 inhibition by rucaparib switched the mechanism of killing induced by MNNG-camptothecin combinations from parthanatotic to apoptotic (Fig. 5e,g). Thus, these data confirm that PARP1 mediates the interaction between parthanatotic and apoptotic death pathways, leading to antagonism and single agent dominance.

## DISCUSSION

A common step in the development and evaluation of new compounds is to define a drug’s mechanism of action, which is generally defined as a drug’s direct binding target. Drug combinations could, in principle, then be designed based on known or perceived interactions between the targeted proteins. Here, we explore a complementary strategy of defining a drug’s mechanism through its mechanism of cell killing, rather than its binding target. Our drug combination screen featuring drugs with varied killing mechanisms reveals a strong enrichment for non-additive drug-drug interactions, and in particular, enrichment for an extreme form of drug antagonism, which we call single agent dominance (SAD). Our statistical and computational analyses reveal that death onset rate is a major determinant of drug dominance. Furthermore, we successfully used drug-induced death rates to predict new SAD phenotypes in previously untested drug combinations. Future studies should determine the extent to which differences in drug activation rate predicts SAD combinations, or other non-additive drug-drug interactions, in other contexts.

The phenotypes uncovered in this study underscore the complexities of cell death regulation and the need for a greater mechanistic understanding of how different forms of cell death are controlled, both regarding the molecular mechanisms and kinetics of activation. Regulated cell death provides an attractive target for therapeutic intervention, as cell killing is required for curative therapies^38^. As we demonstrated in the case of parthanatos and apoptosis, pathway interactions, or “crosstalk”, likely account for the non-additive responses observed in these drug combinations. This is a particularly striking example, as PARP1 inhibitors are being explored in clinical settings, generally in combination with other cytotoxic therapies, and generally without regard to whether these companion therapies induce apoptotic or parthanatotic death^39, 40^. Thus in the case of PARP inhibitors, our data suggest that the effectiveness of these agents should also depend on the mechanisms by which companion compounds kill cells and, more importantly, the relative rates of activation of these drugs. More generally, considering the overall degree of antagonism and SAD within combinations of cell death targeting drugs, our data highlight the existence of widespread inhibitory crosstalk between death pathways. In most cases, however, we simply lack requisite knowledge on the death pathways activated by each drug and the molecular mechanisms by which most forms of cell death are regulated. In the long-term, predicting non-additive drug-drug interactions should be greatly improved by a deeper understanding of the network of interactions between different subtypes of regulated cell death.

A major benefit from our study is the identification of a “rule” that can be used to streamline the evaluation of drug combinations, namely that rates of drug-induced cell death can be used to predict drug-drug antagonisms. Recent studies have highlighted that drug synergy is not needed, and generally not observed, for clinically efficacious drug combinations^6, 7^. Nonetheless, drug-drug antagonism is likely to hinder treatment efficacy, particularly strong forms of antagonism, such as single agent dominance. Since drug-drug interactions are difficult to predict, current strategies rely mostly on screening drug combinations. This process is laborious due to the combinatorial expansion of possible drug combinations, which is further complicated by the fact that drug-drug interactions often depend on the doses used, the order in which the drugs are applied, the environment in which the drugs are applied, and the genotype(s) under evaluation^10, 41–44^. Our study suggests that drugs may not need to be tested in combination in order to avoid SAD combinations, if the rates of drug induced death onset are known or can be measured (Supplementary Fig. 8).

Thus, generating more detailed measurements of single drugs, while modestly more laborious in some experimental assays, would generate a much greater advantage of effectively removing the combinatorial complexity associated with compound screening (Supplementary Fig. 8). It has previously been recognized that the order of drug addition strongly alters the efficacy of drug combinations^5, 45^; however, it had not been made clear from prior studies how one could predict optimal times in which to stagger the application of drugs. Our study suggests that staggering drug addition or drug release based on the onset of drug-induced cell killing may be an optimal strategy.

Currently, standard approaches do not typically evaluate drug activation kinetics, instead focusing on the relationship between efficacy/potency and dose. These “dose-response” relationships have been the central focus of drug pharmacology data for over a century, and these relationships clearly reveal important insights about a given drug. The kinetic features of a drug are generally not predictable from single time point dose response data. Our study reveals that these “rate-response” relationships are observable in kinetic data, and that these relationships also produce unique insights into the nature of a given drug or drug combination. Given the complementarity of pharmacological and kinetic data, evaluation of both of these types of data should become a new standard.

## METHODS

### Cell lines and reagents

U2OS cells were obtained from ATCC and maintained at low passage numbers (generally less than 20 passages from the original vial). Cells were grown in Dulbecco’s modified eagles medium (DMEM) supplemented with 10% FBS, 2 mM glutamine, and penicillin/streptomycin. Drugs were purchased from either Selleck Chemicals, APEXBio Technology, or Sigma-Aldrich. SYTOX green was purchased from Thermo-Fisher. Poly/Mono ADP Ribose Rabbit mAb (83732) was purchased from Cell Signaling Technologies. Purified rabbit anti-active caspase 3 (559565) and Alexa Fluor® mouse anti-cleaved PARP (558710) were purchased from BD Pharmingen. Secondary antibodies were purchased from LICOR Biosciences and BD Biosciences.

### U2OS mKate^2+^ and BAX/BAK^-/-^ cell-line generation

NucLight red mKate2+ U2OS cells were generated by spin-fecting 6.33×106 TU/mL of NucLight Red virus (Essen Biosciences) with 8 ug/mL Polybrene and 1.5×10^6^ cells at 2,000 rpm for 2 hours at 37°C. After spinning, fresh DMEM was added and the cells were incubated overnight. On the next day, cells were replated onto a 10 cm dish and grown to confluence. mKate+ cells were selected by FACS. BAX/BAK-/- U2OS cell line was generated using the pX330-puro plasmid with a hSpCas9 and BAX (GACAGGGGCCCTTTTGCTTC) or BAK (AGACCTGAAAAATGGCTTCG) sgRNA insert. U2OS cells were transiently transfected using FuGENE HD transfection reagent (Promega). BAX/BAK-/- cells were selected for using the BH3 mimetic Navitoclax (10 µM) for 5 days. Following Navitoclax treatment, cells were clonally selected.

### SYTOX based measurement of cell death kinetics

SYTOX green was tested for linearity by plating cells at a 2-fold cell dilution from 20,000 cells to 1 cell in 5 µM SYTOX. After adhering, cells were lysed with 0.1% triton in PBS for 1.5 hours. Fluorescence was measured on a Tecan M1000 plate reader at 503/524 excitation/emission. Gain was set to capture linearity and for fluorescence to be at ∼85% saturation of the detector. For drug-induced cell death measurements, cells were seeded at 1500 cells per well of 384-well optically black plates, or 2500 cells per well of 96-well optically black plates, and adhered overnight on the day prior to drug addition (“day −1”). On day 0, drugs were prepared at 5X final concentration (10X for 96-well experiments) in DMEM containing 25 µM SYTOX green (50 µM in 96 well experiments). 7-point half log dilutions were prepared in 96 well plates. Drug dilutions were added using an Integra ViaFlo96 electronic pipettor (Integra). Fluorescence readings (504ex/523em) were taken at 0, 4, 8, 12, 16, 20, 24, 30, and 48 hours on a Tecan M1000 plate reader. Additionally, an untreated plate was lysed at t = 0 for initial cell counts by adding 0.1% triton in PBS and incubating 1.5 hours at 37°C at 5% CO_2_. After the 48 hour reading, all experimental wells were lysed and a final fluorescence reading was taken. Lethal fraction and percent viability was calculated using the pre- and post-triton permeabilized fluorescence readings.

### Cell-TiterGlo based measurement of relative viability

Cells were seeded at 2500 cells per well in 96-well plates and adhered overnight. On the day of drug addition 10X drug solutions were prepared as above in DMEM and added to the wells. 48 hours post-treatment, cells were lysed using according to the Cell-TiterGlo (Promega) manufacturer instructions and luminescence was read on a Tecan Spark I plate reader. Relative viability was calculated as percent of DMSO control wells on the same plate.

### Modeling drug induced cell death kinetics

The total number of dead cells at given time points can be determined by measuring SYTOX fluorescence. To determine the LF at a given time point, the starting number of plated cells (t = 0) and the post-triton cell number (t = 48) was fitted to an exponential equation for every treated well (assuming exponential growth)^46^. From the growth curve, the total number of cells could be determined for each condition at any time between 0 and 48 hours. Together with dead cell counts, a LF was determined for each drug and dose at all time points measured. Kinetic LFs could modeled using a Lag Exponential Death (LED) equation previously described previous^21^.

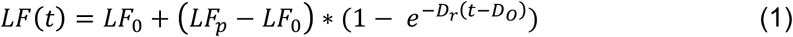

where LF_0_ is the lethal fraction at time 0, LF_p_ is the plateau, D_O_ is the onset time of death, and D_R_ is maximum rate of death. The model was constrained by 0< D_0_ < 48, LF_p_ < 1, and Dr < 2. LF0 was left unconstrained due to basal levels of cell death. Drugs that did not produce max LF values 2x the LF_0_ value were fit to a linear model.

### Flow cytometry based analysis of apoptosis

Cells were seeded at 200,000 cells per well of 6 well dishes on the day prior to treatment and adhered overnight. For time course experiments, drugs were added in a manner such that all samples were collected at the same time. Post treatment, cells were collected and fixed with 4% formaldehyde for 15 minutes. After two washes with PBS, cells were exposed to 100% methanol at −20°C for >2 hours. Cells were washed with PBS and incubated with the active caspase-3 antibody (1:500 dilution, BD Biosciences) in a 50/50 (v/v) PBS-T:Odyssey blocking buffer solution (LICOR). Cells were washed with PBS-T and incubated with the Alexa-fluor 647 cleaved PARP primary and Alexa-fluor 488 goat anti-rabbit secondary antibodies (1:500 dilution, BD Bioscience) overnight at room temperature. FACS samples were run on an LSR II machine with excitation lasers of 488 and 640 nm.

### Live cell image capture

mKate2 expressing U2OS cells were seeded at 200,000 cells per well of 6 well dishes and adhered O/N. On day 0, drugs were added to wells in addition to 50 nM SYTOX. Images were captured using an EVOS FL Auto microscope with a 10X objective using GFP (470/510) and Texas Red (585/624) light cubes (Life Technologies).

### Western blot analysis of PARylation

Cells were seeded at 200,000 cells per well in 6-well dishes and adhered overnight. All drugs were added at t = 0 and cell lysates were prepared at indicated time points. Briefly, media was removed from the well and washed 2x with 2 mL of PBS. Cells were lysed by adding 400 uL of SDS-Lysis buffer (50 mM Tris-HCl, 2% SDS, 5% glycerol, 5mM EDTA, 1 mM NaF, 10 mM β-GP, 1 mM PMSF, 1mM Na_3_VO_4_, protease inhibitor and phosphatase inhibitor tablet). Lysates were spin-filtered through 0.2 um multi-well filters (Pall). After filtration, lysate concentration was determined by the Pierce BCA Protein assay kit according to the manufacturer’s instructions (Thermo). Lysate concentrations were normalized to 0.5 mg/mL for SDS-PAGE loading. Samples were run on pre-cast 48-well gels and transferred using a semi-dry fast transfer (i-BLOT, Invitrogen). Membranes were blocked in a 50/50 (v/v) PBS-T/Odyssey blocking buffer solution for 1 hour at room temperature, incubated overnight at 4°C in primary antibody, and then stained with secondary antibodies conjugated to infrared dyes (LICOR). Blots were visualized using a LICOR Odyssey CLx scanner.

### Data analysis and statistics

All statistical analyses and curve fitting was performed using MATLAB software. PCA was performed in MATLAB after z-scoring data. Analysis of flow cytometry data was performed using FlowJo and Western blot analysis was done using LICOR Image Studio. Combination indices were calculated by Bliss independence using normalized area over the curve values (AOC). Observed AOC values were divided by the expected given: (A+B)-(A*B).

### Data availability

Source data for the drug combination screen are included in Supplementary Table 1. All other data, MATLAB analysis scripts, will be made available upon request.

## ACKNOWLEDGEMENTS

We thank current and past members of the Lee lab and all members of PSB for their helpful comments and critiques during the execution of this study. In addition, we thank Marian Walhout, Job Dekker, Amir Mitchell, and Justin Pritchard for their thoughtful comments during the preparation of this manuscript. The px330-puro-hSpCas9 plasmid was a kind gift from Thomas Fazzio’s lab. This work was supported by National Institute of General Medical Sciences of the National Institutes of Health (R01GM127559 to MJL), and the American Cancer Society (RSG-17-011-01 to MJL). RR, BDL, and PCG were supported by a NIH training grant (Translational Cancer Biology Training Grant, T32-CA130807).

## AUTHOR CONTRIBUTIONS

This project was conceived by RR and MJL. Combinatorial drug screen was designed, executed, and analyzed by RR. BDL, PCG, and MJL helped with the execution of the combination drug screen. Imaging experiments and STACK analysis were performed by RR and HRS. Drug evaluation and annotation of drug mechanism of action were performed by RR, AJJ, PCG, and MSS. Flow cytometry based analyses were performed and analyzed by RR and MEH. All other statistical analysis and modeling was conducted by RR and MJL. Manuscript was written and edited by RR and MJL.

## CONFLICTS OF INTEREST

The authors report no conflicts of interest.

## SUPPLEMENTARY INFORMATION

**Supplementary Figure 1.**
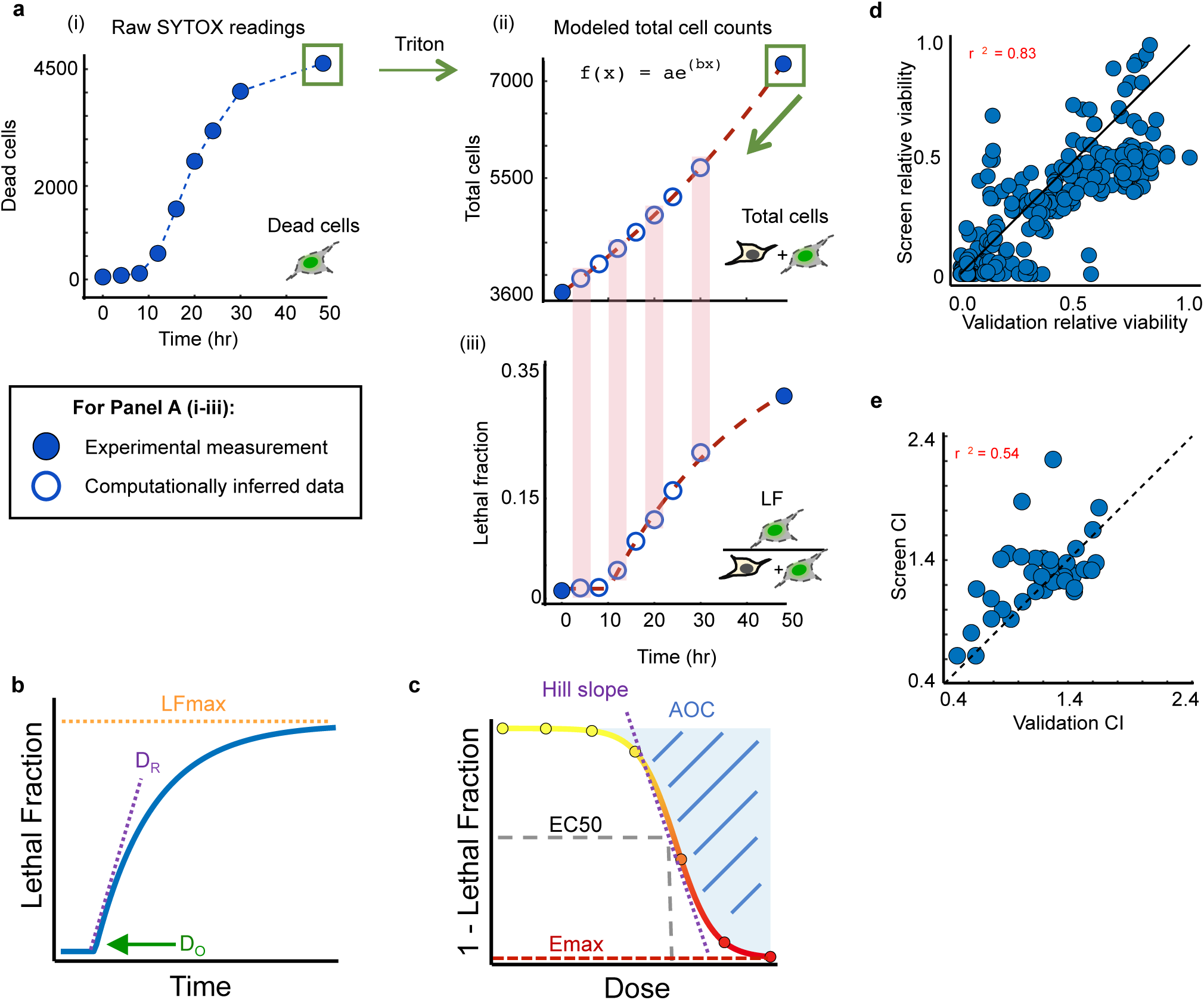
Estimation of lethal fraction dynamics over time using a “pseudo-STACK” plate reader based SYTOX assay. **(a)** Modeling LF over time. (i) Raw SYTOX readings are measured at specific time points (dead cell counts). (ii) After triton permeabilization total cell counts can be determined at 0 hour and 48 hour time points (filled blue circles). Total cell counts can be modeled to an exponential fit (red dashed line) to calculate total cell counts at measured time points (unfilled blue circles). (iii) Modeled total cell counts were used to find the LF at time points where SYTOX measurements were taken (unfilled blue circles) and fit with the LED equation (red dashed line). **(b-c)** Definitions of the kinetic (b) and pharmacometrics (c) used in this study. **(d)** Comparison of combination indices from the cell death screen measured by SYTOX and from combination indices measured using Cell-titerGlo in validation experiments. **(e)** Comparison of relative viability between the cell-death screen and validation experiments as measured in (b).

**Supplementary Figure 2.**
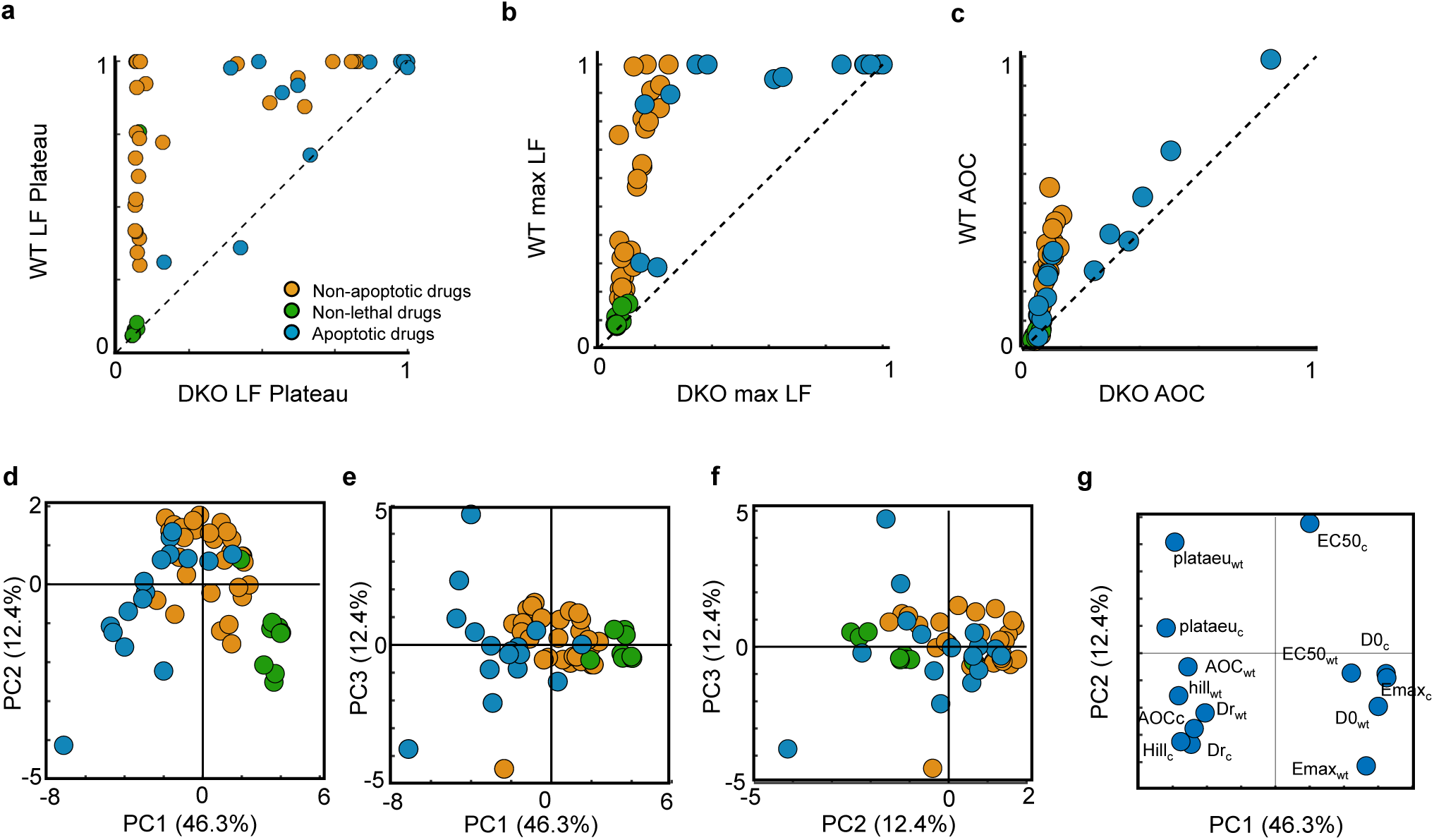
Evaluation of drug mechanism of killing. (a-c) Drug class determination by kinetic and pharmacological response comparison in WT and BAX/BAK^-/-^ (DKO) U2OS cells. Correlation of responses for LED plateau (a), maximum LF (b), and pharmacological AOC (c) between WT and DKO cells. Green – non-lethal compounds; teal – non-apoptotic compounds; Yellow – apoptotic compounds. **(d-f)** Principal component analysis of pharmacological and kinetic parameters in WT and DKO U2OS cells. Projection of observations on PC 1 and 2 (d), PC 1 and 3 (e), and PC 2 and 3 (f). Drugs colored as in (a). **(g)** Loadings plot for PC’s 1 and 2. Loadings correspond to responses from wild-type (wt) or the BAX/BAK^-/-^ clone as in (c).

**Supplementary Figure 3:**
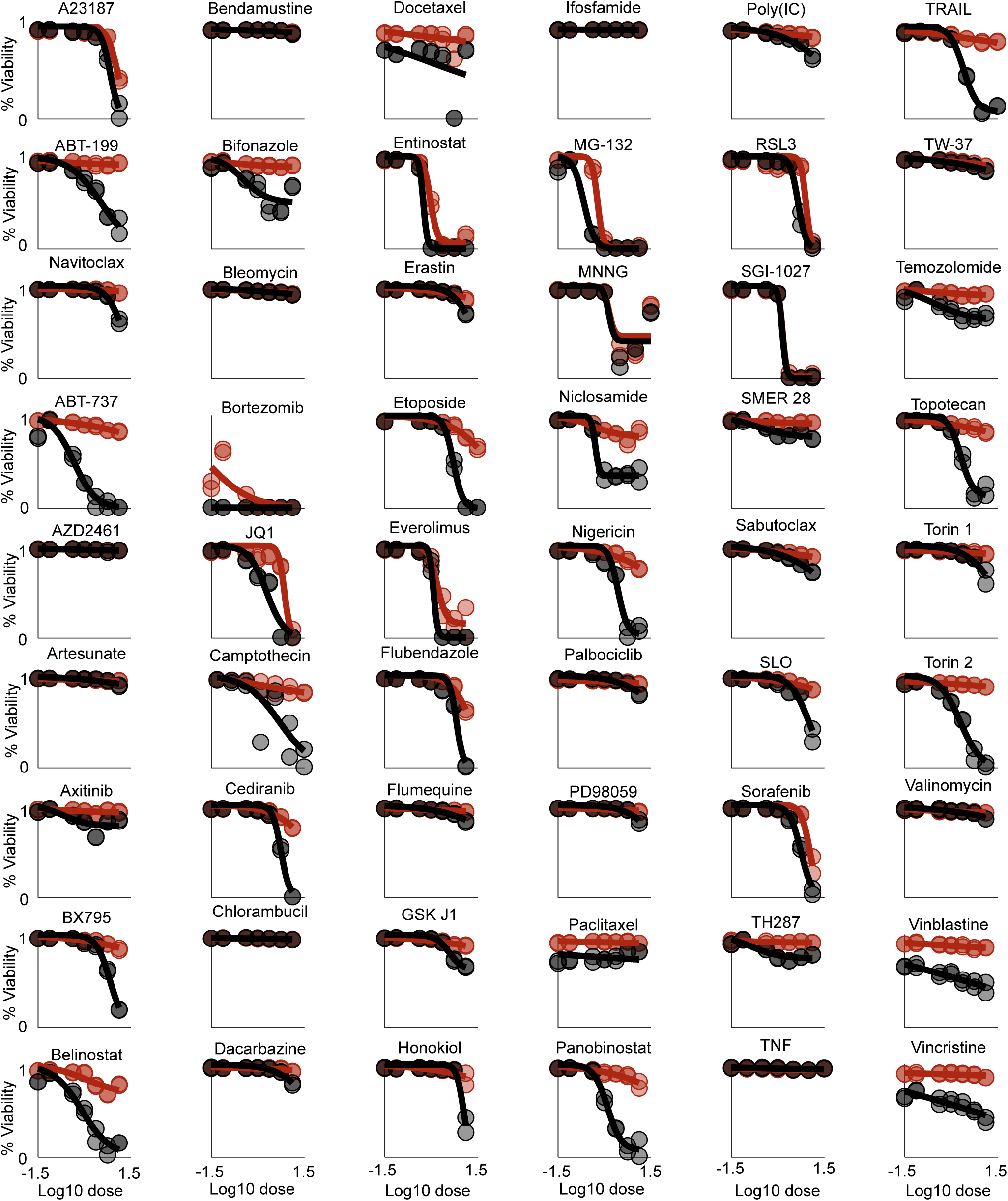
Pharmacological evaluation of drug response. Relative cell viability of the 54 drugs used in the cell death screen in WT U2OS (black) and BAX/BAK-/- DKO cells.

**Supplementary Figure 4.**
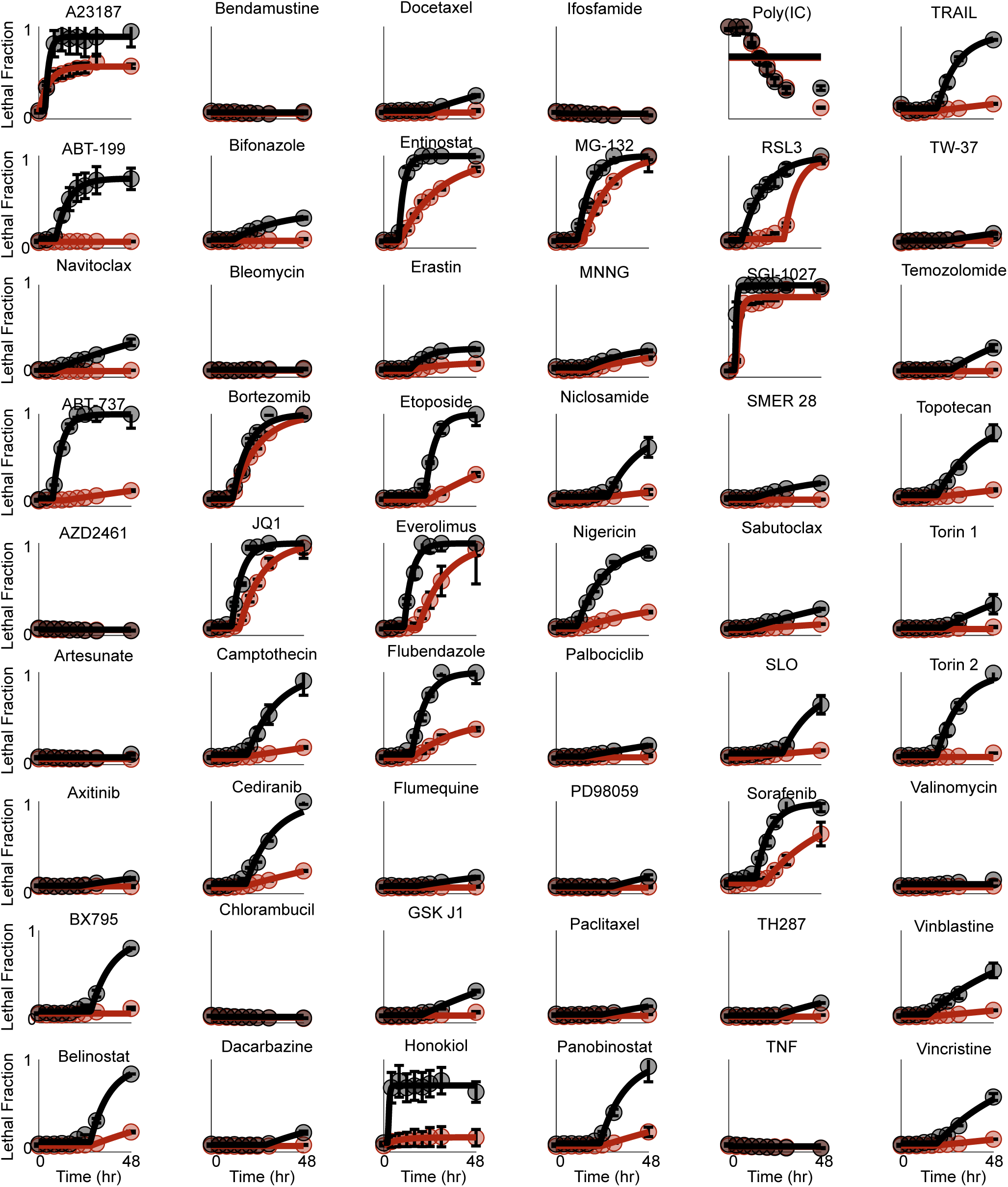
Kinetic evaluation of drug response. . Lethal fraction kinetics in WT U2OS (black) and BAX/BAK^-/-^ DKO cell lines. Data points are mean +/- SD. Solid lines are LED model fits.

**Supplementary Figure 5.**
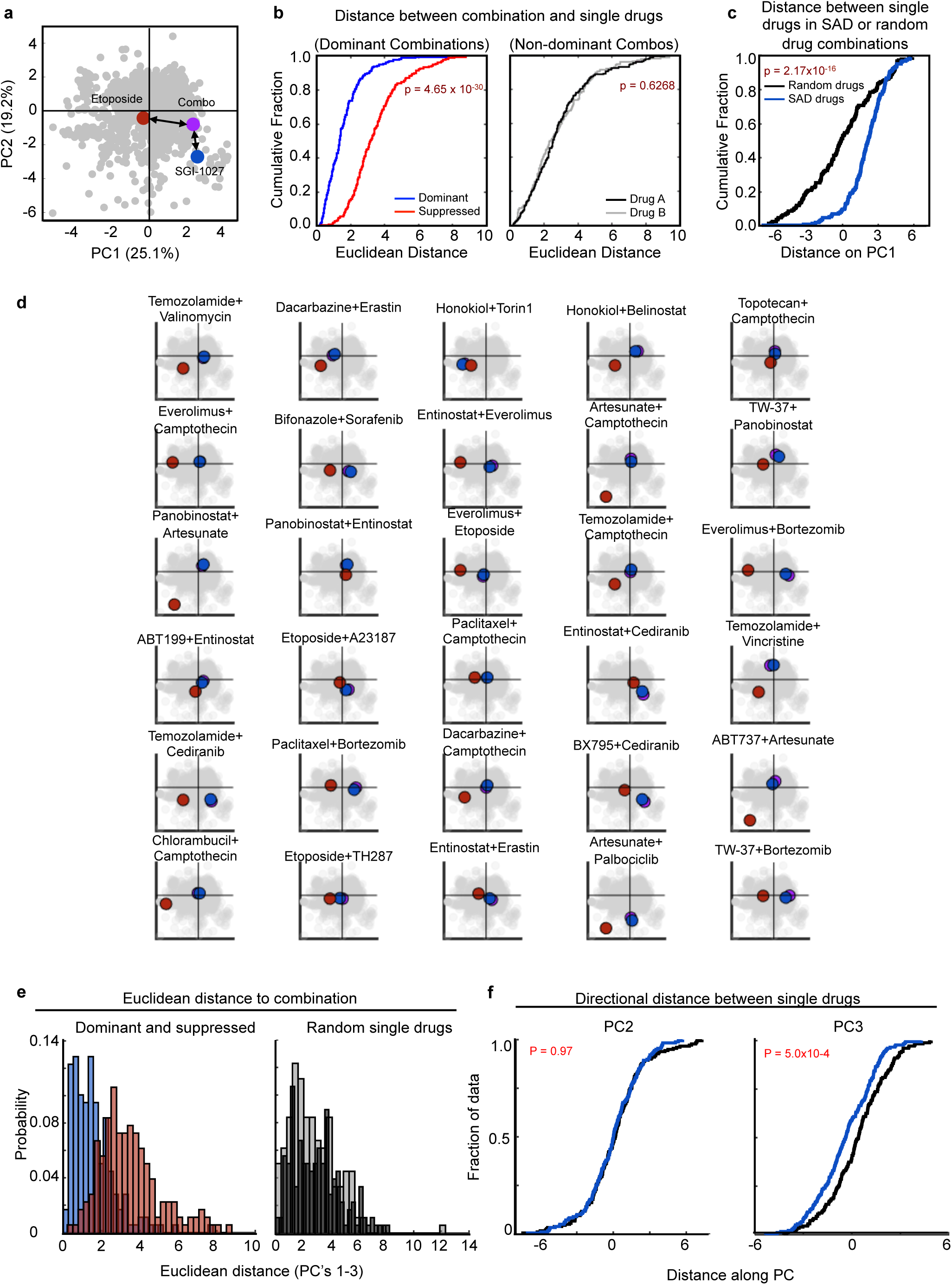
Statistical validation of drug dominance. **(a-c)** Dominant drugs project closer to the drug combination than the suppressed drug in PCA space. **(a)** Example of the relative distances between a dominant (SGI-1027) or suppressed (etoposide) drug and the observed SAD combination. All other combinations and single drug projections are colored grey. **(b)** The Euclidean distance between single drugs and their combination determined along PC’s 1-3 for SAD combinations (left) or non-SAD combinations (right). **(c)** The distance between dominant and suppressed drugs, or random drug combinations on PC1. p-values for panels B and C generated using the KS test. **(d)** Spatial relationship between dominant drug (blue), suppressed drug (red), and SAD combination (purple) on PC’s 1 and 2 for 25 of the 179 SAD combinations. **(e)** Histograms of Euclidean distance between single drugs and corresponding combinations. (left) Distribution of distances between SAD combinations and the dominant (blue) or suppressed (red) single agents. (right) Distribution of distances between random combinations and corresponding singles. **(f)** ECDF of distances on PC2 (left) and PC3 (right) between dominant and suppressed drugs that make up SAD combinations. p-values were determined by a KS-test.

**Supplementary Figure 6.**
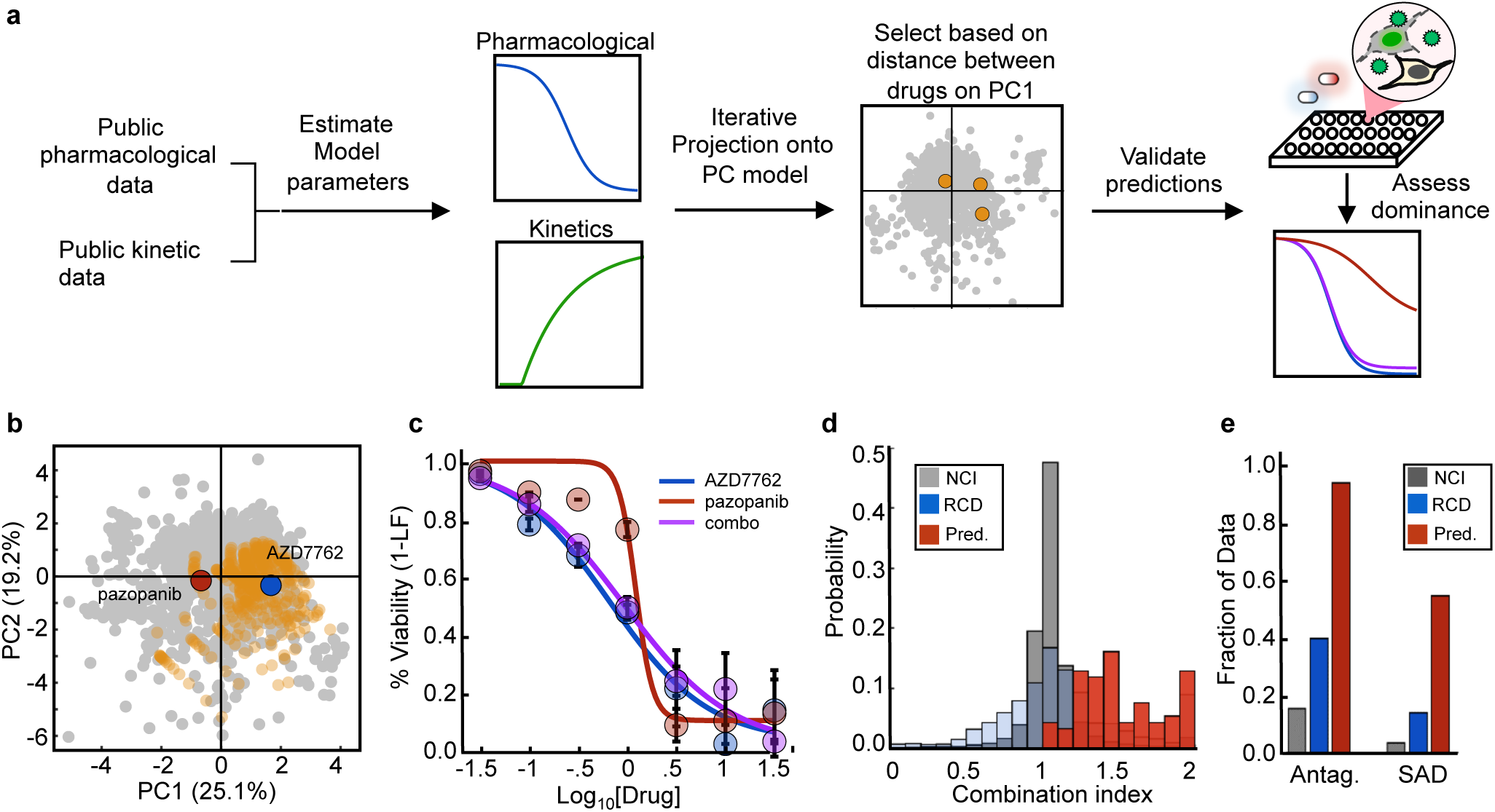
Drug activation rate based classification accurately predicts SAD combinations. **(a)** Workflow to predict new SAD combinations. Public data were used to estimate model parameters for pharmacological and kinetic data and single drugs were projected iteratively onto the PCA model. Based on directional distance on PC1, drug combinations were classified as being SAD or non-SAD. Combinations were validated for antagonism and SAD using SYTOX. **(b)** Example of an estimated drug projection onto the PCA model. Publically available data were used to create a probabilistic array of projections for each single drug (red and blue). All probabilistic projections are shown in orange. A single predicted SAD combination shown, with AZD7762 predicted to dominate pazopanib. **(c)** Validation of a predicted SAD combination. Percent viability was measured by SYTOX and assessed for antagonism and SAD. Blue – AZD (predicted dominant); Red – pazopanib (predicted suppressed); purple – combination. Data are mean +/- SD. **(d)** Histogram of combination indices for all 77 combinations in the model validation set. Distribution of NCI ALAMANAC data (NCI, grey), cell death screen (RCD, blue), and predicted SAD combinations (Pred., red). **(e)** Summary statistics from predictive modeling. Percentages of antagonistic and SAD combinations. Colors correspond to those in (d).

**Supplementary Figure 7.**
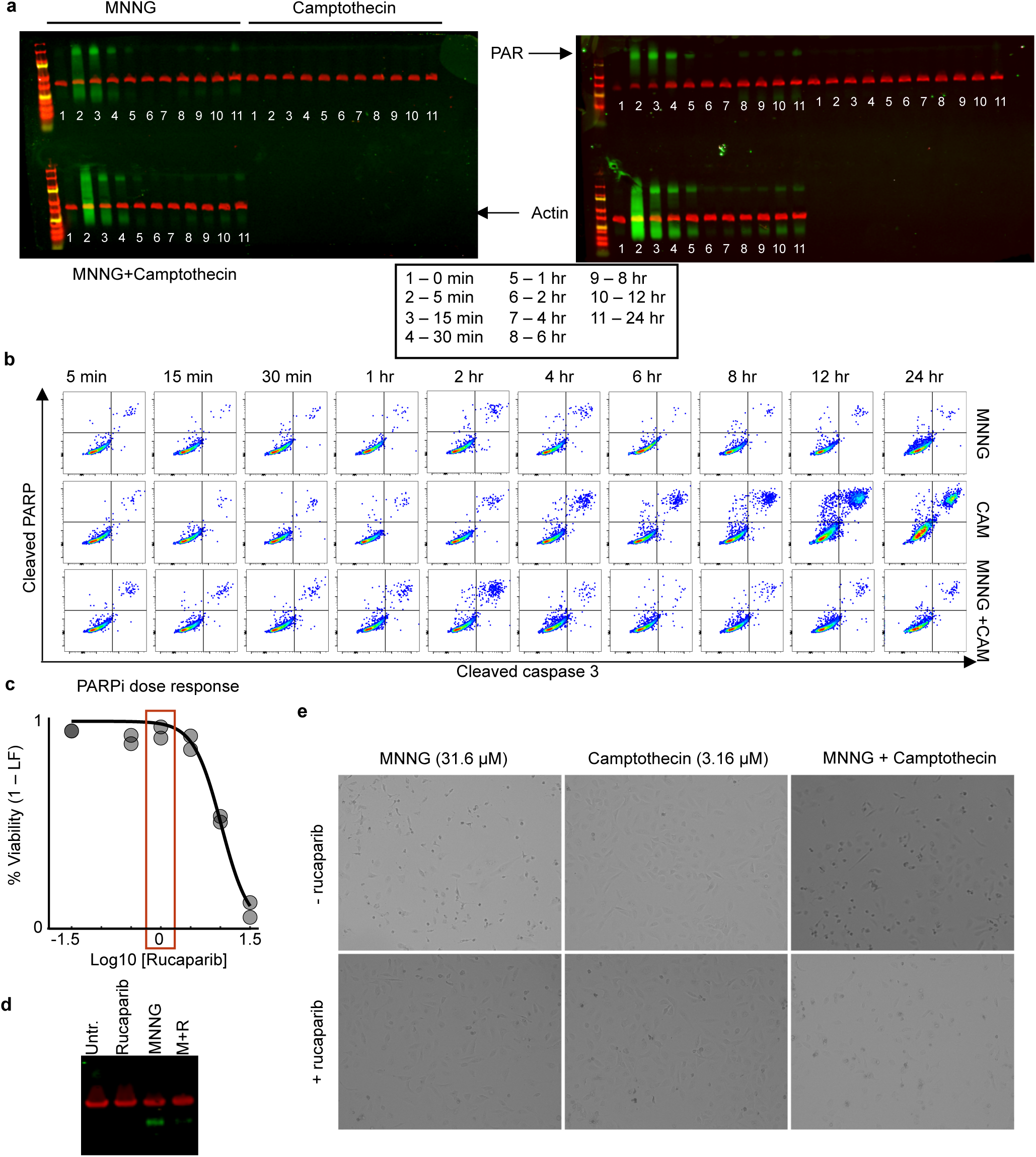
Validation of parthanatotic death induced by MNNG. **(a)** Western blot of protein PARylation. Time course of PAR levels following treatment with MNNG, Camptothecin, or MNNG+Camptopthecin. Blots are biological replicates and time points are identical for each treatment condition. Green – PARylation; Red – β-actin. **(b)** Cleaved PARP activity over time. Representative FACS plots used to quantify cleaved PARP over time (See also Fig. 5b). **(c)** Rucaparib dose response. Rucaparib efficacy was quantified by SYTOX. A sub-lethal dose (1 µM, highlighted red) was chosen for subsequent experiments for PARP inhibition. **(d)** PAR activity in the presence of PARP inhibition. U2OS cells were treated with 1 µM rucaparib, 100 µM MNNG, or the combination for 24 hours. **(e)** Parthanatotic cell morphology. Trans illumination images of U2OS cells following indicated treatments +/- 1 µM Rucaparib at 48 hours.

**Supplementary Figure 8.**
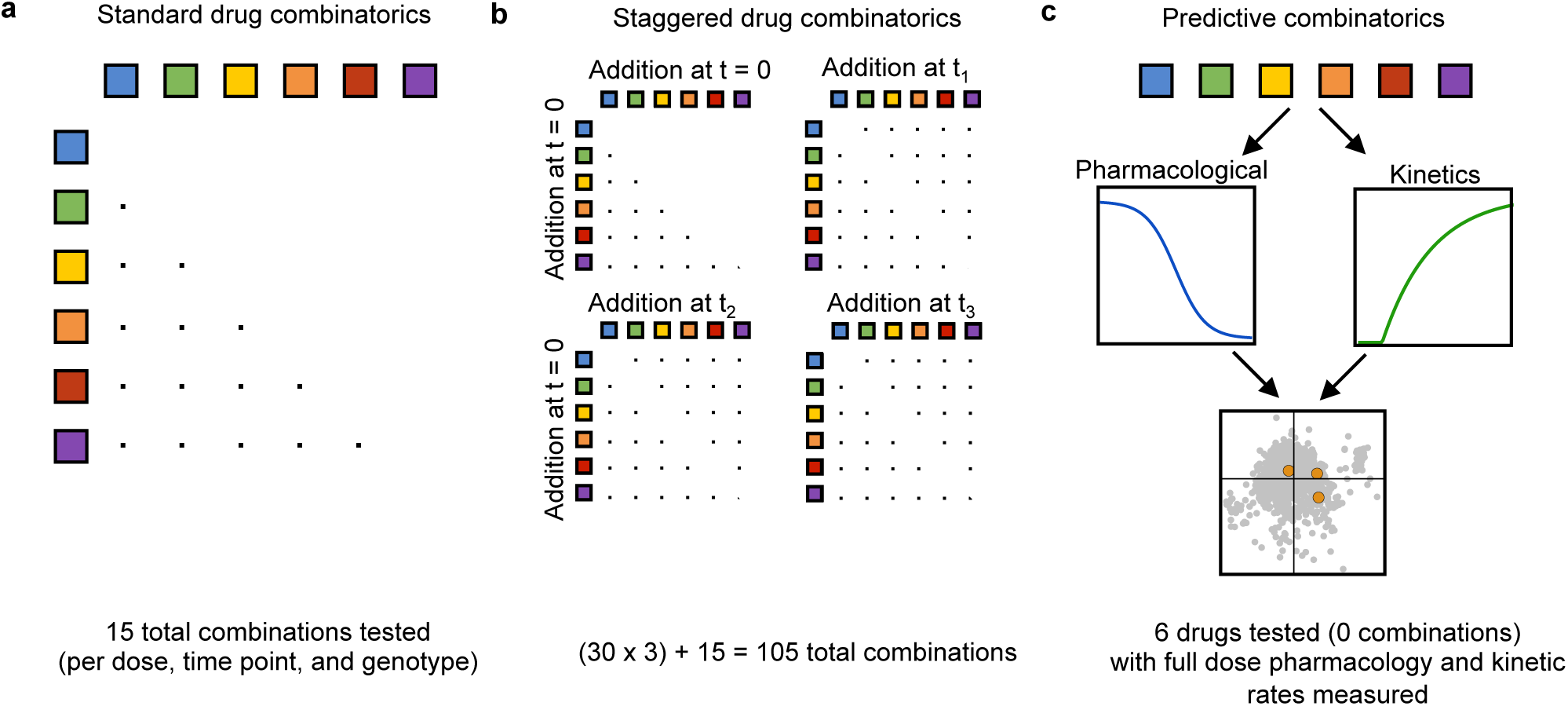
Rate based evaluation of single drugs may improve issues related to combinatorial expansion associated with testing drugs in combination. **(a)** Drug combinations when drugs are added at the same time. Example of testing all pairwise combinations of drugs, given that the order of drug addition does not matter. These data are for a single dose, evaluated at a single time point, in a single genetic background. **(b)** Drug combinations when relative timing of drug addition needs to be controlled. In example shown, drug “A” is added at t = 0 and drug “B” is added at time t_i_. Total combinations shown for 3 temporal staggers, again at a single dose and time point, in a single genetic background. **(c)** Predictive combinations using a rate based classifier. Using predictive models, testing drugs in combination may not be necessary. For instance, to identifying and avoiding SAD combinations, drugs can be tested only individually. Single drug kinetics and pharmacological parameters are then measured and modeled using PCA to predict antagonistic interactions, and the optimal drug temporal regimen needed to avoid SAD combinations in favor of additivity.

**Supplementary Table 1 : Combination drug screen data.** Full drug response, including pharmacological parameters, kinetic rates, and combination indices for all drugs and drug combinations tested in Fig. 2.

Supplementary Table 2: **PCA scores for all drugs and drug combinations**

**Supplementary Code: Analysis script for computational back-fitting of lethal fraction over time**

